# Comparative ChIP-seq (Comp-ChIP-seq): a novel computational methodology for genome-wide analysis

**DOI:** 10.1101/532622

**Authors:** Enrique Blanco, Luciano Di Croce, Sergi Aranda

## Abstract

In order to evaluate cell-and disease-specific changes in the interacting strength of chromatin targets, ChIP-seq signal across multiple conditions must undergo robust normalization. However, this is not possible using the standard ChIP-seq scheme, which lacks a reference for the control of biological and experimental variabilities. While several studies have recently proposed different solutions to circumvent this problem, substantial technical and analytical differences among methodologies could hamper the experimental reproducibility and the quantitative accuracy. Here we propose a local regression strategy to accurately normalize ChIP-seq data in a genome-wide manner. Overall, our proposed experimental and computational standard for comparative ChIP-seq (Comp-ChIP-seq) will increase experimental reproducibility, thereby reducing this major confounding factor in interpreting ChIP-seq results.

## BACKGROUND

Chromatin is the macromolecular complex of DNA and histone proteins that packs the genome into its basic structural units of nucleosomes [1]. Within chromatin, a plethora of interacting proteins organize the 3D distribution of the genome, regulate multiple gene expression programs, and coordinate the appropriate transmission of genetic and epigenetic information to cellular progeny [1-5]. Alterations in the functionality of the proteins associated with chromatin are intimately linked to severe developmental diseases and cancer [6]. Indeed, one recent comprehensive analysis of 10,437 cancer exomes has revealed that nearly 40% of all cancer driver genes are well-characterized chromatin associated factors, which include transcription factors, chromatin modifiers, chromatin remodelers, and guardians of genomic stability [7]. Due to its biological and pathological relevance, research on chromatin and epigenetics has been a rapidly moving field over the last decade, assisted by the development of novel methods for the high-throughput molecular analysis of the genome.

The development of chromatin immunoprecipitation (ChIP) coupled with the next-generation sequencing (seq) methodologies has been pivotal for characterizing the genomic distribution of a vast collection of chromatin-associated proteins, histone post-translational modifications (PTMs), and histone variants [8-11]. The striking impact of the ChIP-seq technology is based in its relative technical simplicity (which allows it to be adopted by most experimental laboratories), its sensitivity and accuracy for mapping the genomic distribution of proteins, and the standardization of the experimental and computational methods to efficiently analyze such a volume of information. Fifteen years ago, the Encyclopedia of DNA Elements (ENCODE) project was launched as a collaborative initiative to catalog the complete set of functional elements in the human genome in selected cell lines [12, 13]. More recently, the International Human Epigenome Consortium (IHEC) has generated a comprehensive high-resolution reference map for the epigenome of major primary human cell types [14, 15]. The ChIP-seq method has been central for building the cartography of functional elements of the human genome in both of these international collaborative efforts.

Current research efforts aim to identify the changes in the occupancy profiles of chromatin-bound factors or histone modifications in two or more cell types, metabolic states, and/or pathological situations. However, in its traditional scheme, ChIP-seq is essentially a semi-quantitative method that enables the researcher to determine the relative occupancy of one factor in a given genomic region, with respect to the rest of the genome. In other words, there is no direct correlation between the amount of ChIP signal in the output and the biological activity of the binding element in different scenarios [16]. Several sources of biological and technical variability can be overlooked, thereby hampering any direct comparison of ChIP signal strength between different conditions [17]. For instance, the method of preparation of the sequencing material by ChIP is a source of technical variability. An apparent increase in genomic occupancy of a chromatin factor could simply be the result of variability in the efficiency of immunoprecipitation or DNA elution between experiments. Moreover, while running the sequencer, a standard practice is to mix equal proportions of barcoded libraries to run the samples in a multiplexed manner. Therefore, even a substantial global reduction of a histone variant occupancy per cell would remain hidden in a ChIP-seq experiment after normalizing by total number of reads [17]. Although the consistent replication of ChIP-seq experiments can reveal the biological tendency in the interacting strength of the chromatin factor, a robust normalizing strategy is required to accurately compare ChIP-seq results across experimental conditions.

To overcome the influence of technical variabilities in the biological interpretation of ChIP-seq, several groups have reported different strategies based on the use of internal reference controls (spike-in), which provides a feasible solution to accurately normalize comparative ChIP-seq (Table 1). Originally developed to correct gene expression measurement in microarrays and RNA-seq experiments [18, 19], the spike-in strategy is based on combining the experimental sample with an amount of exogenous material (either from another species or synthetically produced) that is constant between experiments. Both the experimental sample and the spike-in are processed and analyzed in parallel. As long as the amount of spike-in ChIP signal is constant, the observable differences in the experimental samples across conditions can be exclusively attributed to biological variation. Eventual differences in the spike-in signal can be computationally equilibrated to eliminate technical variability, and the same correction is then used to normalized the experimental signal.

**TABLE 1:** Table summarizing the different methodologies developed to normalized ChIP-seq signal with internal controls. For each spike-in method we initially provide the following basic information: bibliographical reference, type of material used to normalize and the method used to capture the reference control. Next, we assess the strengths and weaknesses of each method according to several parameters: cell counting requirements, the constrain in epitope conservation, the capacity to estimate the average ChIP signal per cell, the possibility to monitor the fragmentation efficiency in the reference material and the stability of the ChIP-seq signal in the reference control. Finally, we indicate the computational method undertaken for each analysis.

| Name | ChIP-Rx | Single-antibody ChIP | Two-antibodies ChIP | Parallel-factor ChIP |
| --- | --- | --- | --- | --- |
| Reference | *Orlando, D.A. et al, 2014* | *Bonhoure, N et al. 2014* | *Egan, B. et al, 2016* | *Guertin, M.J. et al, 2018* |
| Normalizing sample | Xenogenic cells | Xenogenic chromatin | Xenogenic chromatin | Parallel ChIP with sample material |
| Capturing method of spike-in material | Antigen conservation between sample and spike-in material | Antigen conservation between sample and spike-in material | Specific antibody for spike-in material (H2Av) | Second antibody for endogenous protein |
| Cell counting | Required | Not required | Not required | Not required |
| Epitope conservation | Required | Required | Not required | Not required |
| Estimation average ChIP signal per cells | Possible | Not possible | Not possible | Not possible |
| Monitoring fragmentation on spike-in material | Not possible | Possible | Possible | Possible |
| ChIP -seq profiling of spikein material | Constant | Constant | Constant | Assumed to be constant |
| Computational analysis | Absolute fold-change correction | Signal fold-change correction | Absolute fold-change correction | Linear local correction |

Current strategies for ChIP-seq normalization using spike-in diverge in: *i*) the type of biological material to be used as a reference sample for normalization; *ii*) the capturing method of the spike-in material; and, *iii*) the computational method used to analyze the sequencing data. It is important to mention that the use of different strategies can introduce inconsistencies to the final results, as each method presents its own benefits and limitations. In addition, the existence of several alternatives might complicate decision-making for researchers about when and how to apply a normalization method for comparative ChIP-seq. With the aim of providing a practical, unified reference framework for comparative ChIP-seq analysis, we have now evaluated the benefits and limitations of the experimental methodologies and analytical pipelines currently available. We have proposed an experimental best practice and, in addition, we have designed a companion bioinformatics pipeline for analyzing comparative ChIP-seq data in a genome-wide manner. If widely used, we believe that this computational pipeline can serve as a reference for increasing data reproducibility between laboratories, thus overcoming one of the major drawbacks of using ChIP-seq data.

## RESULTS AND DISCUSSION

### Comparison of current experimental methodologies

Over the last five years, up to four alternative strategies have been proposed based on the spike-in concept to deal with the problem of a lack of comparability between multiple ChIP-seq samples. An additional normalization method has been also developed using nucleosomes reconstituted from recombinant and semisynthetic histones on barcoded DNA prior to immunoprecipitation (ICeChIP, Internal Standard Calibrated ChIP) [20]. Although considerably useful for the accurate quantification of local histone modification densities, we have focused on protocols compatible with cross-linked ChIP-seq against any potential target of interest. We have summarized the principal benefits and limitations of each technique in Table 1.

In 2014, two different laboratories independently pioneered the development of a similar strategy for comparative ChIP-seq normalization, by introducing xenogenic material from a second different species into the experimental model [21, 22]. The rationale behind both approaches is that the ChIP-seq signal obtained from a fixed amount of spike-in material can be used as an internal reference to normalize the experimental ChIP signal across different samples. Thus, Guenther and collaborators mixed *Drosophila melanogaster* S2 cells with human cells before cell lysis [21], while Delorenzi and colleagues added a constant amount of fragmented chromatin from human cells into previously-fragmented mouse chromatin [22]. Despite the conceptual similarities, the initial decision on which spike-in material should be added to the experimental sample raises important technical considerations that can influence the resulting biological interpretations. The addition of xenogenic cells of the spike-in material enables the whole procedure to be monitored from the beginning, thereby minimizing the impact of technical variabilities for the biological interpretation. Further, by mixing cells, it is possible to tackle eventual changes in genomic ploidy (*e.g.* due to genomic instability, or differences in cell cycle progression), thereby providing an estimation of average ChIP-seq signal per cell. This quantitative estimation is not possible when mixing fragmented chromatin. However, the option of mixing cells is only available when the number of cells can be evaluated accurately (*e.g*. cells growing on a dish), or when the experimental sample and spike-in material are fragmented with the same settings. On the contrary, when number of cells in the sample is uncertain (*e.g*. animal tissue samples), or when the experimental sample and spike-in material require different settings for fragmentation, the addition of the fragmented spike-in material at chromatin level is a more appropriate option.

In the previous strategies, the sample and the spike-in material are captured using the same antibody, limiting both strategies to using antigens that are highly conserved between both classes of material (for instance, histone modifications). To circumvent this problem, Trojer and colleagues introduced a smart solution by using a second antibody for a fly-specific histone variant (H2Av) to capture the spike-in material [23]. This strategy aims to avoid the cross-reactivity constraint of the experimental antibody and to reduce any potential variability due to competition between the spike-in control and the experimental material, which usually exceeds the amount of spike-in material by far.

In order to overcome both epitope conservation restrictions and the use of a xenogenic spike material, a fourth normalizing strategy has been recently proposed: rather than using exogenous material, Guertin and co-workers recommend including a second antibody against an endogenous target present in the experimental chromatin, as an internal control [24]. This second antibody is used to profile the genomic occupancy of a pervasive chromatin factor (*e.g*. CTCF) whose genomic distribution is assumed to be unchanged between different cell types and/or treatments and which is clearly distinguishable from the experimental target [24]. Nonetheless, although this method avoids preparation of xenogenic material, we consider that it still makes two important assumptions, which could not be always true: i) that the reference endogenous factor remains stably associated under different experimental conditions; and, ii) that the genomic distributions of endogenous reference target and the experimental target do not overlap.

### Towards an experimental framework for comparative ChIP-seq experiments

Considering the benefits and limitations of the different strategies (reviewed in Table 1), we propose the addition of exogenous xenogenic fly cells, whenever possible, and the use of a second antibody against a fly-specific histone variant, as a best practice for comparative ChIP-seq normalization for mammalian genomes. We recommend using fly material as the spike-in because: 1) the genome sequence has been extensively assembled; 2) the fly chromatin has been largely characterized at epigenetic levels; 3) the evolutionary distance between fly and mammalian genomes is sufficient to allow an unambiguous alignment of the reads [21, 23]; and 4) fly cells are relatively easy to culture with standard tissue culture procedures and instruments. Moreover, the genomic occupancy profile of the fly-specific H2Av is already characterized [23], which can be extremely useful as an additional control point for assessing ChIP-seq performance.

Taking into account the previous considerations, we have designed a practical guide under the form of a decision tree to systematically implement a consistent protocol for the addition of the spike-in material when performing ChIP-seq experiments. In our roadmap, we state some key questions that are relevant for deciding which type of spike-in to add (*e.g.* fly cells or fragmented chromatin, see data flow diagram in Fig. 1). These guidelines take into consideration that: (*i*) spike-in material should be present in all samples at equal amounts at the earliest step during the ChIP-seq procedure; and (*ii*) spike-in material should be present in a low-enough quantity (giving a significantly lower number of reads as that of the experimental reads) to not interfere with the actual ChIPseq experiment yet still give an accurate normalization in the final sequencing data. As previously reported, the number of spike-in reads in the final sequencing step should be at least one million reads, and approximately, 2%–5% of the experimental genome, to minimize the changes in overall material used for ChIP-seq [21, 23]. This final amount of reads can be influenced by the ratio of the mixture as well as by the quality of the antibody and/or the abundance of the target in the experimental condition. Taking into account these considerations, and the relative ratio between the size of the fly genome and the two most widely used mammalian experimental models (mouse and human), we recommend the use of different final mixtures (Fig. 1).

**Figure 1:**
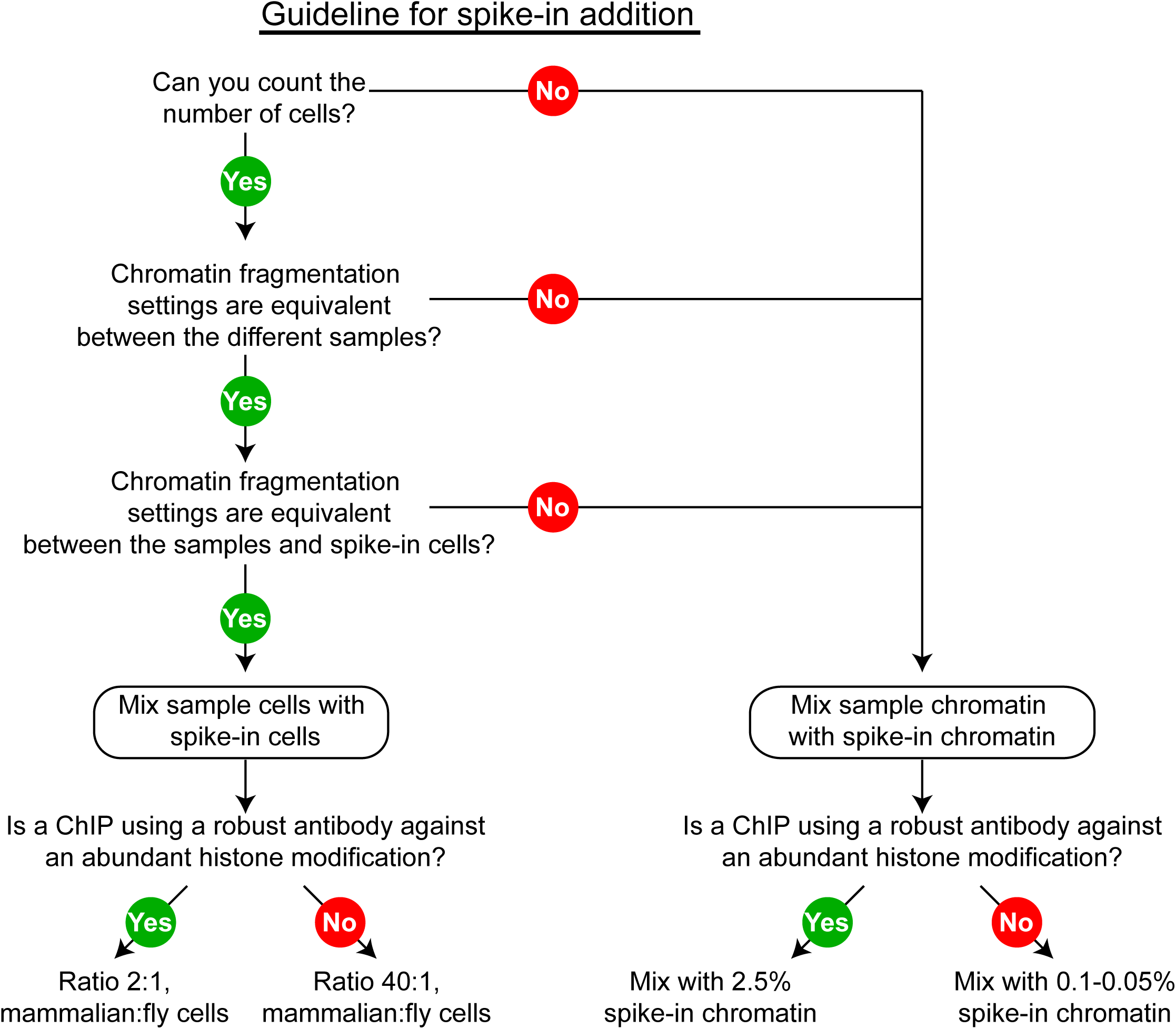
Flow chart to select the most appropriated strategy for a Comp-ChIP-seq experiment.

### Comparison of current computational methodologies

Traditionally, the standard ChIP-seq normalization method uses the total number of mapped reads per million (RPM) to correct for possible bias introduced by the differences in the sequencing depth among samples. With the recent spike-in methods (see Table 1), the mapped reads from the spike-in control are employed to correct the experimental ChIP-seq signal strength.

At the computational level, the rationale of the different spike-in methodologies is the same: as long as a constant amount of spike-in material is added to the experimental samples, and as long as the samples and the spike-in material are processed and analyzed together, the correction factor computed to eliminate the differences in the spike-in signal can be used to normalize the experimental sample signal. However, the different methodologies differ in the computational approach to correct the spike-in and, consequently, the experimental sample.

Several authors have proposed to take advantage of the number of mapped reads of the spike-in sample (*e.g*. *Drosophila melanogaster*) to correct the overall ChIP-seq signal from the main organism studied (*e.g*. *human*) [21, 23]. Although initially appealing, this fold-change correction presents in our opinion important shortcomings, such as: *i*) the spike-in reads mapped not only along the ChIP-seq peaks but also over background regions are used for computing the correction factor; and *ii*) the correction factor is uniformly applied to all experimental reads in the actual experiment, treating both non-specific and specific signal loci with the same correction value. Alternatively, the use of a normalization factor computed derived from a pre-defined list of known targets in the spike-in organism was also proposed [22]. This option excludes any confounding background signal, but the correction is only applied to a computational pre-defined loci list with specific ChIP-signal, thereby relying in the accuracy of the selected computational peak caller to define the list of positive loci.

Recently, with the aim of overcoming the limitations of the previous approaches, Guertin et al. proposed to apply a linear local regression method [24]. This approach computes a correction coefficient, defined by a linear regression model, for the systematic and gradual correction of the pre-defined ChIP-seq peaks from the reference. After this, the coefficient is also used to correct a pre-defined subset of peaks in the experimental sample. In addition, such a linear correction method implements a statistical approach to calculate the probability and strength of differentially bound loci. Guertin et al. (23) also showed an increased sensitivity (of about 10%) in the detection of differentially bound target loci as compared to the previous absolute fold-change correction. The conceptual improvements of this approach stem from the fact that the correction factor gradually increases along with the informative power (as number of reads) of the peaks. However, the addition of a computational step to pre-select the real signal loci in this strategy, similar to the non-linear correction method, could introduce an additional bias step, as the consistency in the outcome of available peak calling tools is limited [25]. In addition, this analysis would impede the genome-wide evaluation of the signal-to-noise ratio, thereby limiting the informative power of the ChIP-seq.

### Benchmarking a novel local regression method for comparative ChIP-seq in a genome-wide manner

With the aim to enable a genome-wide comparative analysis of ChIP-seq experiments, we developed a novel computational method that performs the genome-wide normalization of ChIP-seq data adapting the spike-in control correction to the class of genomic region (Fig. 2). Our strategy overcome the limitations from the previous strategies, as: 1) it normalizes ChIP-seq signal over the complete genome, and not just from a subset of selected regions; 2) in order to compute the correction factor, the influence of the reads from background regions, although dominating over the total number of reads, is minimized; and, 3) the correction factor derived from the spike-in material is not uniformly applied over all the experimental ChIP signal, instead, it is increasingly and gradually applied from background to positive ChIP signal regions. Our approach, inspired by the spike-in-based RNA-seq quantification methods described above [18, 19], shares conceptual similarities with the recent linear local correction approach [24] and consists in the application of a local regression, in this case, over all the genome-wide bins determined along the chromosomes (see Methods). Thereby, our method is able to introduce a distinct correction factor to each bin in the genome, depending on its class. First, a local regression (LOESS) is computed from the bins in the spike-in genome in order to accommodate the two ChIP-seq conditions compared into the same best-fit line. Next, the values from the real experiment (experimental reads) are corrected following the previous local normalization calculated using the spike-in bins (Fig. 2). Under this approach, the adjustment on a region containing a true ChIP-seq signal is expected to be substantially higher than the change computed for bins with background signal.

**Figure 2:**
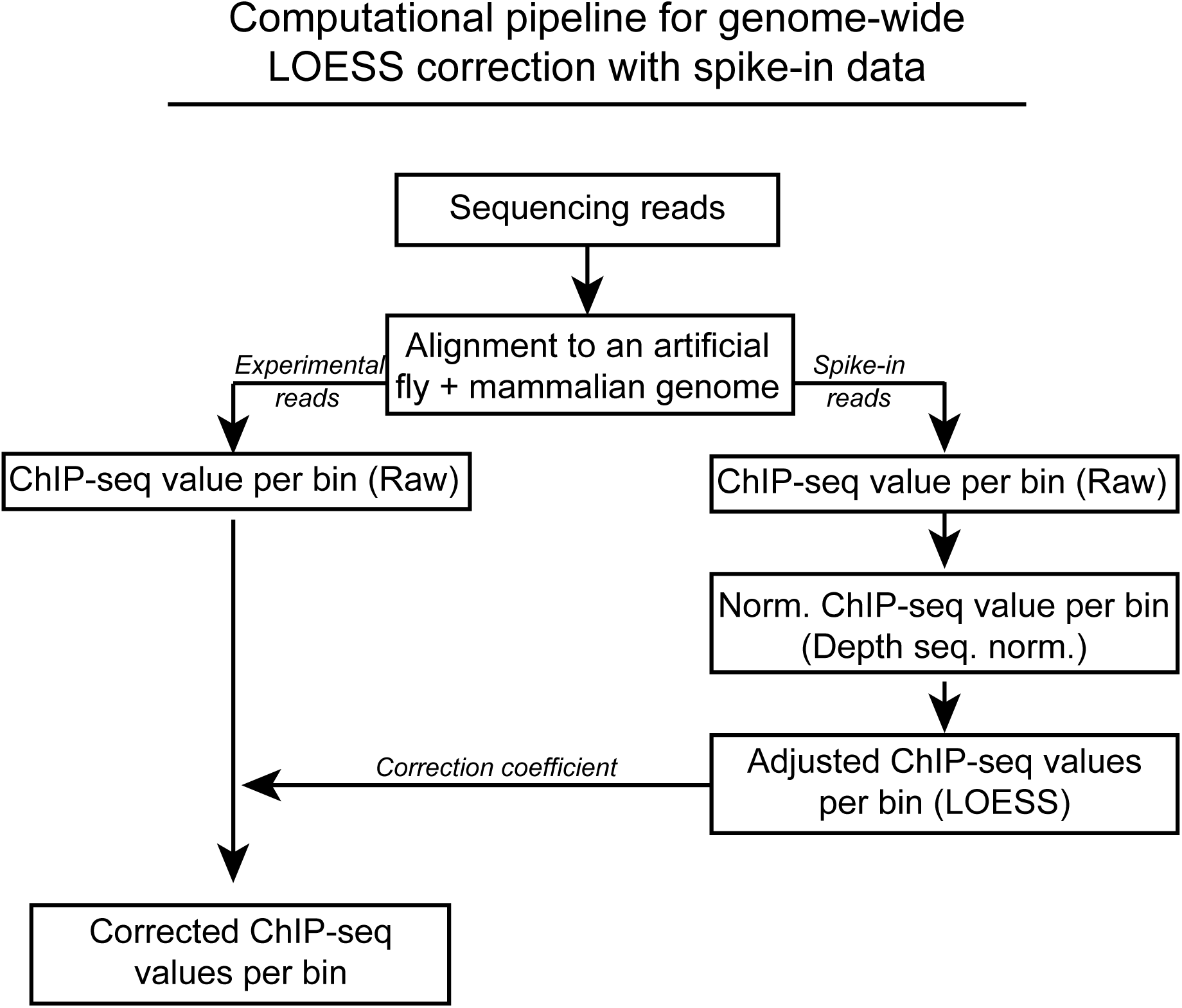
Computational pipeline for normalizing ChIP-seq data using spike-in ChIP-signal in a genome-wide manner. A diagram summarizing the computational analysis protocol (see Methods section for details).

To assess the accuracy of our proposal, we compared the performance of our LOESS method with the fold-change strategy on a reference dataset [21, 23](Fig. 3). We took advantage of the available ChIP-seq data published by Guenther et al. that included fly material as spike-in control [21]. In this study, the authors artificially generated a pre-defined ChIP signal gradient for the di-methylation of lysine-79 histone H3 (H3K79me2). To achieve a controlled range of distinct conditions, they mixed different proportions of Jurkat cells that had been untreated or treated with a selective inhibitor for the H3K79-methyltranferase DOT1L (EPZ5676). The mixture aims to reflect the global change in the average H3K79me2 level per cell. For our benchmarking, we selected two completely different conditions: i) the 25:75 (DMSO:EPZ5676) proportion, which has higher levels of H3K79me2; and, ii) the 75:25 proportion, with lower levels of H3K79me2 (Fig. 3). Finally, a constant amount of fly cells was used as an internal reference control for normalization.

**Figure 3:**
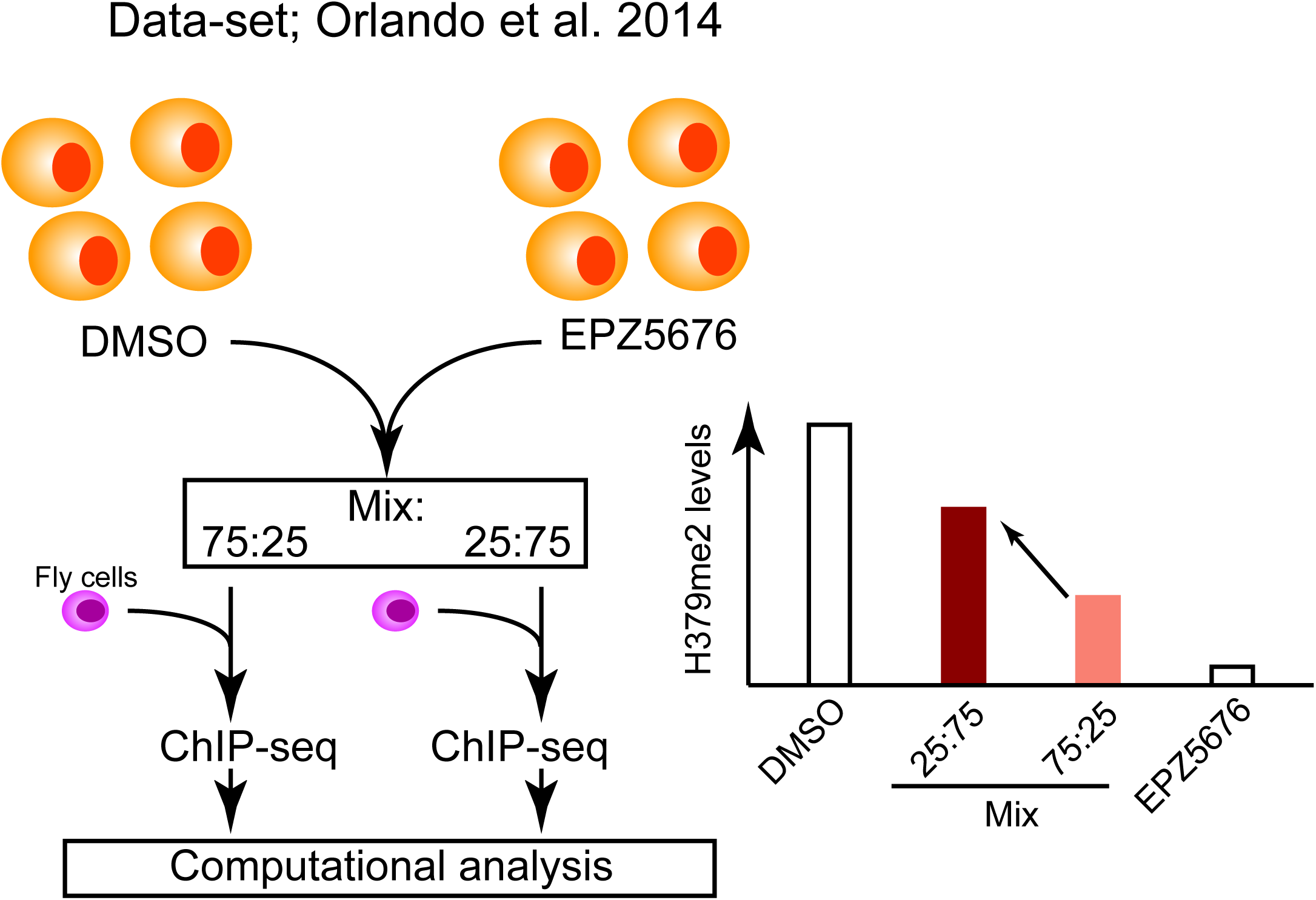
Available datasets used in this study. Scheme representing the experimental approach undertaken in [21] to generate a gradual ChIP-seq signal for H3K79me2.

We processed the H3K79me2 samples from the two different conditions. As indicated in our computational pipeline (Fig. 2), first we mapped the resulting sequencing reads to an artificial genome in which we included the human and the fruit fly chromosomes. Next, we segmented the genomes of the sample (human) and the spike-in control (fly) into bins of 1 Kb. After separating the mapped reads into human and fly, we assigned the average ChIP-seq value of H3K79me2 in both conditions within all bins from both genome segmentations (see Methods). These initial values, which were not corrected by any normalization method, were considered to be the raw value (Fig. 2). We then used the spike-in data to compute the normalization of each bin using the fold-change (FC) methodology or our LOESS approach, respectively. For FC correction, we used the total number of aligned fly reads to correct the experimental human ChIP signal of both conditions, as previously suggested [21, 23]. For LOESS correction, we first normalized the spike-in sample for the total number of reads, and then applied the LOESS correction in the fly bins to the best fit-line (Fig. 2 and 4). Then, we used the same correction factors, depending on the density of reads within the bins, to normalize the human experimental bins. An appropriately analytical normalization using spike-in should display a qualitative and quantitative difference between both experimental ChIP signals in the peaks of H3K79me2, while keeping their background levels equilibrated.

**Figure 4:**
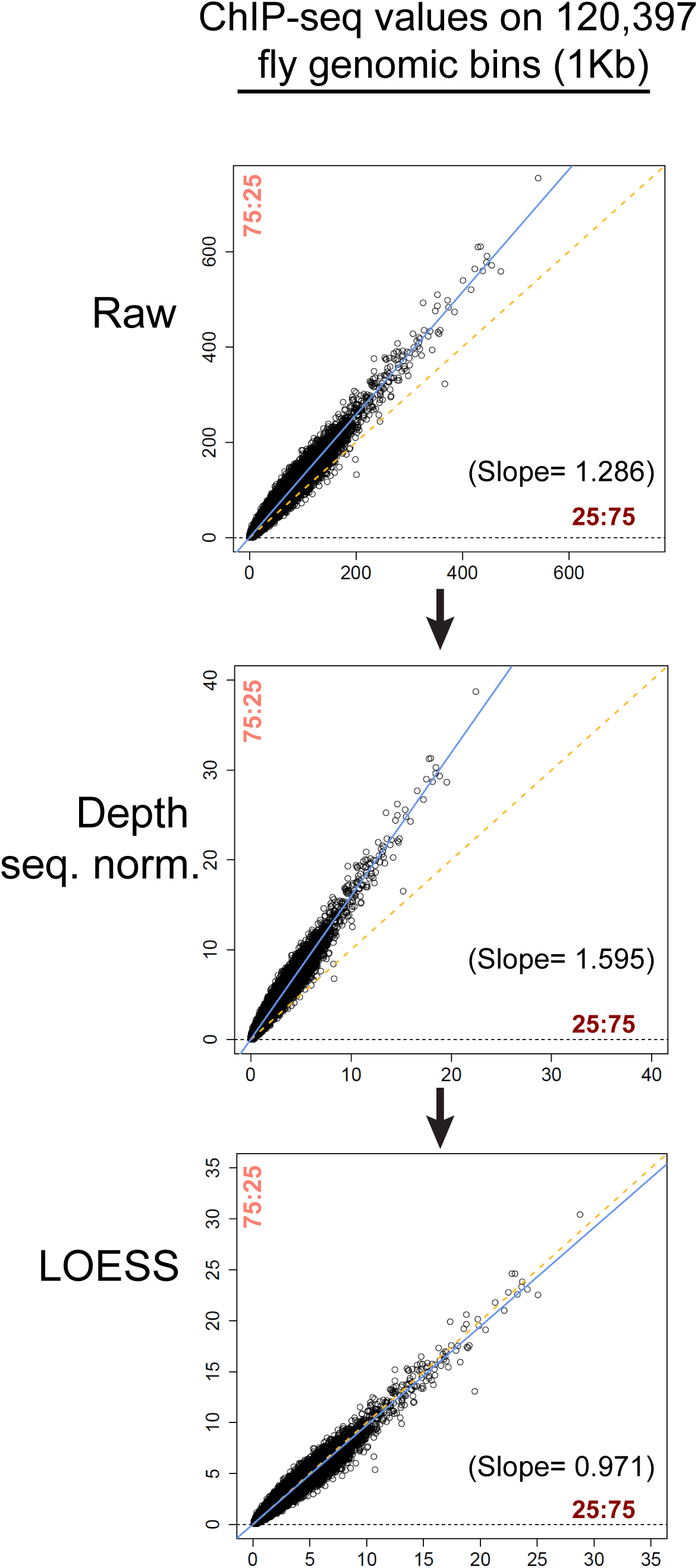
LOESS normalization of the reference genome. Scatterplots showing the distribution of fly bins in both conditions accordingly to the mean value of H3K79me2 on each bin. From top to bottom, we show the raw values, the values adjusted by the sequencing depth and the final values corrected by the resulting local regression by LOESS to the best-fit line.

As shown in Figure 5a, either using the FC normalization or the LOESS approach, ChIP signal strength is remarkably different between the 25:75 and the 75:25 samples. However, after a detailed inspection over the background regions, we consistently found a sustained increase also in such areas when using the FC correction. These genome-wide changes observed in the target occupancy over all background are misleading since the genomic distribution of H3K79me2 must be considered to be equivalent between both samples, as resulting from mixing the same samples with different proportions. Instead, when using LOESS, the ChIP signal at background regions remains equilibrated (Fig. 5a). In order to systematically determine the accuracy of both methods, we compared the average signal of all bins reported as peaks by MACS and the same value for rest of the bins of the human genome (background). As anticipated, meta-plot and heatmap analysis centered on the bins corresponding to H3K79me2 peaks displayed a disproportionate difference on the surrounding bins belonging to the background when using FC correction (Fig. 5b). In contrast, the ChIP signal of background bins remains equilibrated between both conditions when using LOESS correction, as experimentally expected.

**Figure 5:**
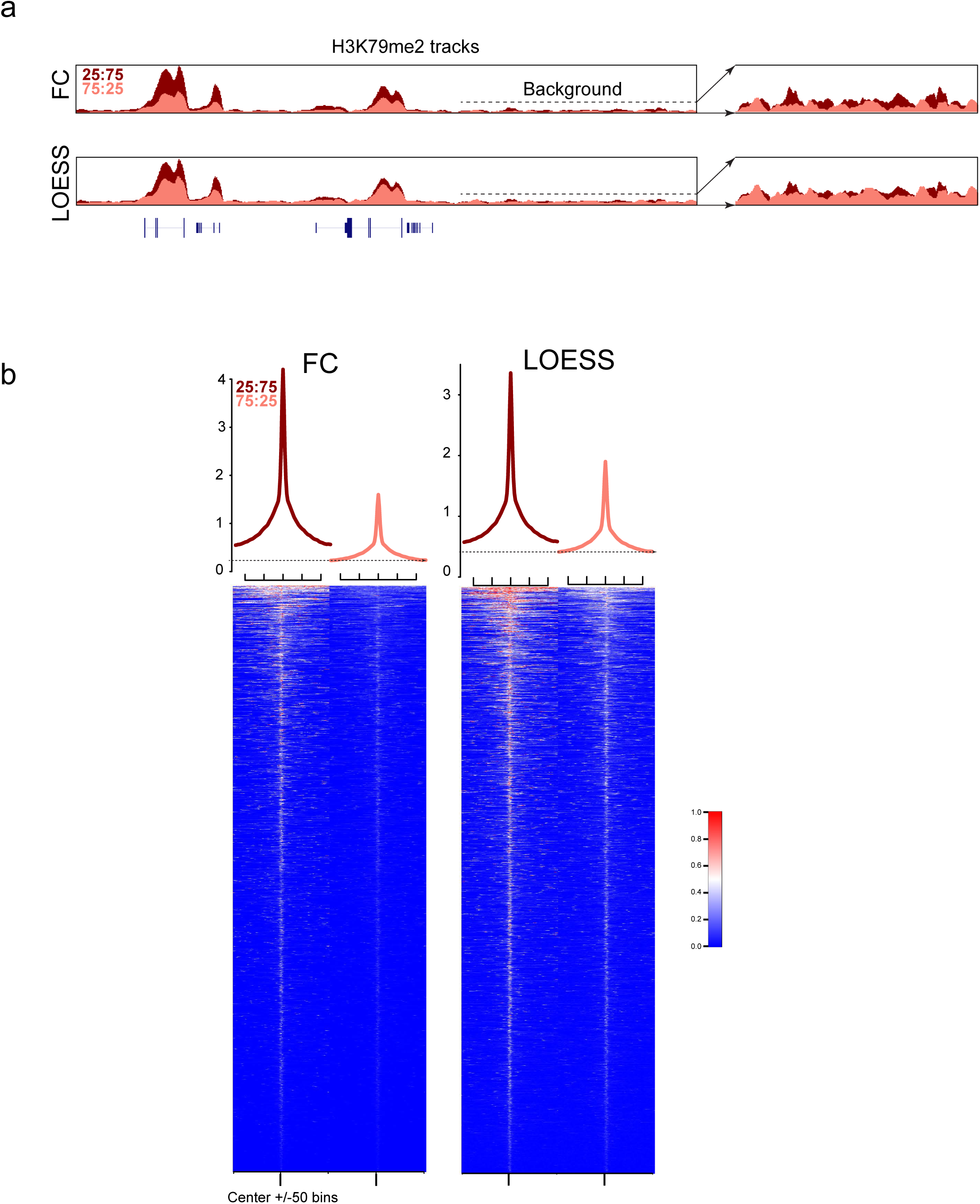
Experimental ChIP-seq signal distribution. **a** Overlayed ChIP-seq tracks from H3K79me2 on 25:75 and 75:25 samples from [21], normalized by FC or LOESS (assembly: hg19, coordinates chr19:23,400,000-23,650,000). On the right, magnification of a background region indicated by the arrows. **b** The metaplots represent the average of ChIP-seq values per bin from a 100 Kb genomic region window centered on the bins containing a peak according to MACS. The heatmaps correspond to the 100 Kb genomic region window from the same bins.

When quantifying the ChIP-seq signal in both classes of bins, the differences in the background normalization resulted even more evident (Fig. 6). While applying the FC correction in human bins in a genome-wide manner, we quantified a general increase in ChIP-signal on the 25:75 sample with respect to the 75:25 sample in the background bins (19.7%, Fig. 6a), which overall constitute more than 97% of total reads (Fig 6b). In contrast, a near perfect balance was strikingly quantified on the bins that constitute the background level of the ChIP-seq from the human experiments when using LOESS correction (+1.7%).

**Figure 6:**
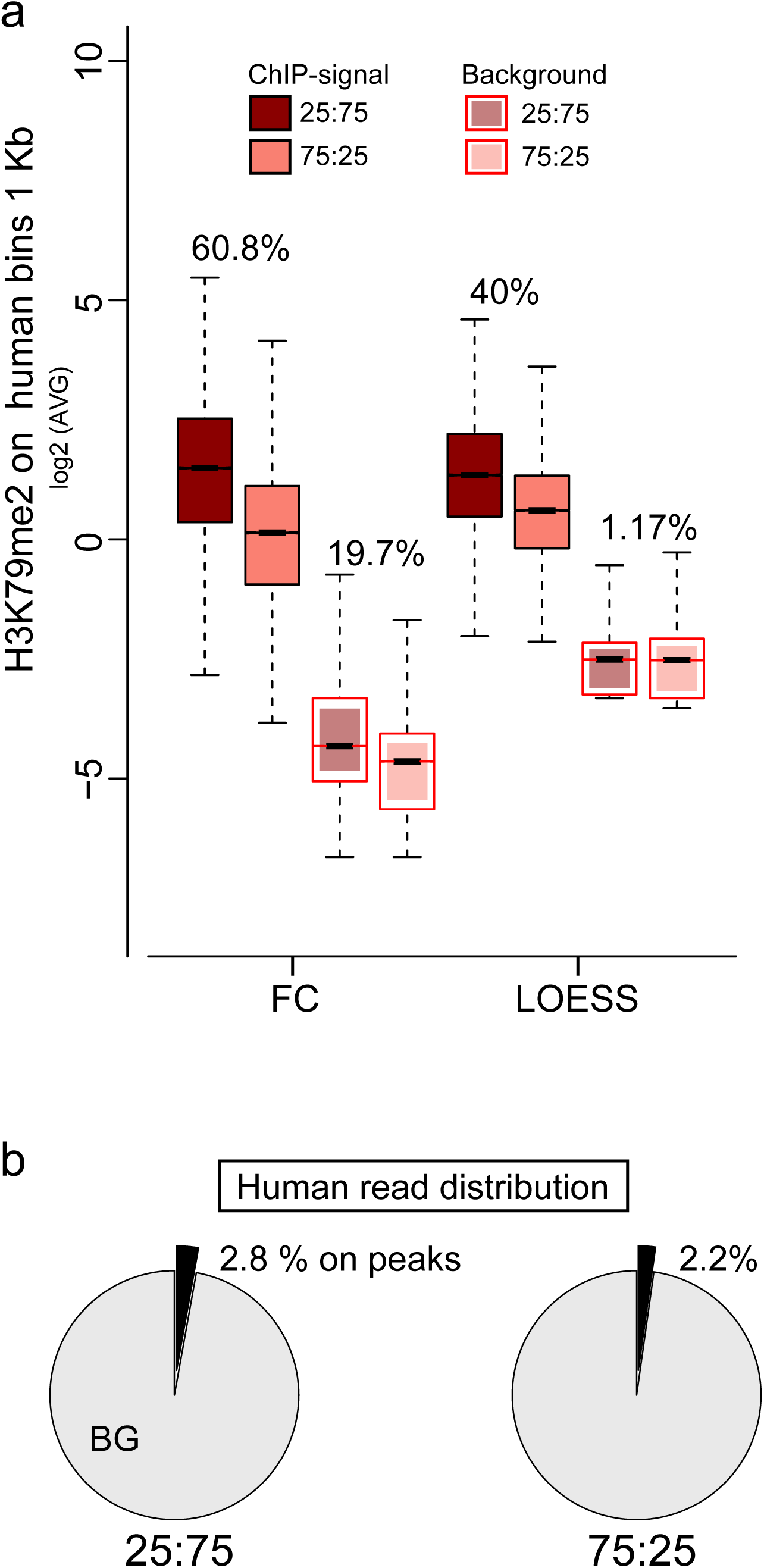
Experimental ChIP-seq signal distribution. **a** Boxplots representing the distribution of H3K79me2 ChIP-seq signal after absolute fold-change normalization (FC), or LOESS normalization, using the spike-in material. The percentage of difference between the medians of boxplots are indicated. **b** Distribution of the experimental human reads in the 25:75 and 75:25 samples between the background regions and ChIP signal regions. Note the low percentage of reads that fall into regions with a peak called.

To our knowledge, the distinct gradual normalization from background to positive ChIP signal regions has not previously been considered in any of the existing normalization techniques. We thus believe our proposal presents a major technical advance for the genome-wide normalization and for comparison of ChIP-seq experiments.

## CONCLUSIONS

The ChIP technique is one of the most widely-used methods in molecular biology [26], since its development over 30 years ago [27-29]. The power of this technique has increased dramatically with the advent of the massive parallel sequencing approaches [8-11]. Indeed, ChIP-seq experiments have become the universal method to delineate the genome-wide maps of distribution of transcription factors, chromatin remodelers, and histone modifications. Since about a decade ago, the original scheme of ChIP-seq has been maintained substantially unmodified from chromatin isolation and fragmentation, immunoprecipitation using specific antibodies, DNA purification from protein complexes, library preparation, and parallel sequencing. The ChIP-seq experiments endow a large proportion of noise, which can be introduced by the crosslinking artifacts, non-specific binding of the antibody, or the high sensitivity of the parallel sequencing techniques. This significant amount of noise results in many cases in most of the reads mapped into regions of the genome that are unrelated with the chromatin target (*e.g.* background zones). A precise determination of the signal-to-noise ratio is, therefore, very relevant for determining the occupancy strength of the targets. Limitations due to potential biases introduced by the technical variabilities in the original ChIP-seq scheme have necessitated the development of strategies to implement an internal reference control across samples for further comparisons. However, there are still potential pitfalls in the application of each approach. Thus, this important problem is still open in many aspects. Very recently, an additional spike-in strategy has been developed [30]. In this new approach, the authors benefit from the genetic diversity of yeast strains to perform an intra-species spike-in using different *Saccharomyces cerevisiae* strains as spike-in and experimental specimen. Beyond its utility in lower eukaryotes, this new method exemplifies the aim to meet an experimental need. Considering the different strategies proposed to normalized ChIP-seq data using spike-in, we provide here a major guideline to plan and perform comparative ChIP-seq experiments, and propose an innovative computational approach for the analysis in a genome-wide manner to fill the gap of the experimental need for an accurate interpretation of ChIP-seq data.

Our proposed standard for the genome-wide comparative ChIP-seq signal has shown to be very effective for the precise comparison of ChIP signals across samples without a pre-defined selection of the loci. This approach is able to correct for possible technical bias and to compute a local correction factor, thereby minimizing the impact of the correction over non-occupied genomic regions. We strongly believe that the systematic use of spike-in references in the ChIP-seq experiments will provide a more precise picture of the dynamics of epigenome in different conditions.

The epigenetic research field is systematically searching to increase the sensitivity for minimizing the amount of cells required for analysis, with the aim of reaching a confident single-cell ChIP-seq. The high variability produced by manipulating single-cell events shall benefit from the introduction of internal reference controls, which are at the conceptual bottom line of the methods reported here. We expect that single-cell Comp-ChIP-seq will bring this technique into a third revolution for tracking the genomic effects of molecules bound to single loci.

## METHODS

### Genome segmentation in bins of 1Kb

Using the UCSC genome browser, for each genome assembly of human and the fruit fly (hg19 and dm3, respectively), we retrieved the list of chromosomes and the corresponding sizes from the following sites:

> http://hgdownload.soe.ucsc.edu/goldenPath/hg19/database/chromInfo.txt.gz
>
> chr1 249250621
>
> (…)
>
> http://hgdownload.soe.ucsc.edu/goldenPath/dm3/database/chromInfo.txt.gz
>
> chr2L 23011544
>
> (…)

Next, we generated each segmentation file of non-overlapping bins of 1 Kb in BED format using the following GAWK command (showing the command for fly only):

> % gawk ’BEGIN{OFS="\t";offset=999;}{for(i=1;i<$2-offset;i=i+offset+1) print $1,i,i+offset;}’ >
>
> fly_1Kb.bed
>
> chr2L    1        1000
>
> chr2L    1001     2000
>
> chr2L    2001     3000
>
> (…)

In total, after filtering the Het and M chromosome files out, we ended up with 3,095,665 bins for the human genome and 120,397 bins for the fruit fly genome.

### Synthesis of the human+fruit fly genome

Firstly, we merged the full set of chromosomes of each genome assembly (hg19 for human and dm3 for *Drosophila melanogaster*) in FASTA format into a single file (genome.fa). Next, we appended the tag “_FLY” to the name of the fly chromosomes in the resulting FASTA file and in the chromInfo.txt file describe above to distinguish human from fly sequences.

>chr1

>chr2

(…)

>chr2L_FLY

>chr2R_FLY

(…)

Finally, we used the command bowtie-build from the BOWTIE suite [31] to generate the corresponding indexes for posterior mapping of the ChIP-seq raw data files.

### ChIP-seq raw data files and mapping

Using the NCBI GEO resource (GEO series accession: GSE60104), we retrieved the raw data of the samples Jurkat_K79_25%_R1 (GSM1465005) and Jurkat_K79_75%_R1 (GSM1465007) from [21]. Next, we utilized BOWTIE [31] to map both FASTQ files of reads over the human+fly genome indexes described above (BOWTIE parameters -p 4 -t -m 1 -S). Finally, we used the SAMTOOLS [32] to filter the unaligned reads (option -F 0×4) out and, by using the “_FLY” tag, the mapped reads corresponding to the fly spike-in control were then separated from the human experimental ones.

### Canonical normalization of the ChIP-seq data values

For each experiment, we separately averaged the number of reads within each human and fly bin using the SAMTOOLS [32]. Next, to assign the final value of one bin depending on the normalization method, we employed the following transformations (formulated below): *i*) absolute values, in millions of reads, which were used as raw values; *ii*) absolute values divided by the total number of human+fly reads per sample, for the traditional normalization; and *iii*) absolute values divided by the total number of fly reads per sample, for the fold-change normalization.

(RAW)

Let X be the number of reads averaged on a particular bin B, the raw value per bin was calculated as:

Raw (B) = X / 10^6^.

(TRADITIONAL NORMALIZATION)

Let X be the number of reads averaged on a particular bin B and N be the total number of human and fly mapped reads, the traditional value per bin was calculated as:

TRADITIONAL (B) = X / N.

(FOLD-CHANGE NORMALIZATION)

Let X be the number of reads averaged on a particular bin B and F be the total number of fly mapped reads, the fold-change value per bin was calculated as:

TRADITIONAL (B) = X / F.

### Calculation of local regression normalization of the ChIP-seq data values

Inspired in a similar treatment proposed for RNA-seq normalization of the RPKMs of spike-in controls [18], we applied the LOESS function from the R library affy to the traditional normalization values, to perform the local regression of data. We instructed the loess function normalize.loess to use the adjustment on the values in the fly spike-in genome as a subset to guide the normalization of the human values. A pseudo-count of 0.1 was added to each value before running the normalization function.

More in detail, we concatenated the fly and the human files of bins containing the traditionally normalized values in both conditions 25:75 and 75:25. The first 120,397 lines of this file corresponded to the fly bins (used as a subset to guide the LOESS) and the rest of the lines to the human bins (normalized in the LOESS method by the corrections to adjust the previous subset of bins). Once the normalization was performed, we separated again the human bins from the fly bins into two different files per ChIP-seq experiment.

### Generation of UCSC tracks, meta-plots and heatmaps

The resulting files of bins and values assigned according to each normalization strategy in both ChIP-seq conditions were directly used to generate BedGraph, meta-plots and heatmaps. In brief, the format of one of these input files was the following (human_bins_LOESS.txt):

> (Column descriptors: BIN ID, normalized value for 25:75, normalized value for 75:25)
>
> chr1*1*1000 0.100055353317942 0.0999446773050048
>
> chr1*100000000*100001000 0.175377416129052 0.173340448679151
>
> chr1*10000000*10001000 0.382830859965691 0.309274962866397
>
> (…)

And the GAWK commands to generate the corresponding BedGraph profiles are:

> gawk ’BEGIN{FS="*"}{print $1,$2,$3}’ human_bins_LOESS.txt | gawk ’BEGIN{OFS="\t"; print “track type=bedGraph name=FG_1Kb_loess_25_human visibility=full color=100,0,0"}{print $1,$2,$3,$4}’ > 1Kb_loess_25_human.bg;
>
> gawk ’BEGIN{FS="*"}{print $1,$2,$3}’ human_bins_LOESS.txt | gawk ’BEGIN{OFS="\t"; print “track type=bedGraph name=FG_1Kb_loess_75_human visibility=full color=200,0,0"}{print $1,$2,$3,$5}’ > 1Kb_loess_75_human.bg;

### Discrimination of bins associated to peaks and to background

MACS [33] was used to identify the list of ChIP-seq peaks along both genomes in the 25:75 and 75:25 conditions. To distinguish between bins that contain ChIP-seq peaks and bins that constitute the background, we calculated the overlap between MACS peaks and the coordinates of the segmentation bins at human and fly chromosomes. We used boxplots to represent the distribution of normalized values that belongs to each class of bins.

#### LIST OF ABBREVIATIONS

ChIP-seq: chromatin immunoprecipitation coupled with the next-generation sequencing
RPM: reads per million
LOESS: locally estimated scatterplot smoothing
FC: Fold-change

## DECLARATIONS

### Ethics approval and consent to participate

Not applicable

### Consent for publication

Not applicable

### Availability of data and materials

The datasets used in this study are deposited in Gene Expression Omnibus (GEO) repository under the accession number GSE60104.

### Authors’ contributions

E.B. conceived and performed the bioinformatics analysis of deep sequencing data and contributed to writing the manuscript. L.D.C. contributed to data analysis and interpretation and to writing the manuscript. S.A. conceived and planned this project, perform data analysis and interpretation, and wrote the manuscript with input from the coauthors.

## Acknowledgements

We specially thank Dr. Cecilia Ballare and Dr. Pedro Vizan, as well as all the members of the Di Croce laboratory for critical reading of the manuscript and insightful discussions and V.A. Raker for scientific editing.

